# Genome-wide analysis of genetic predisposition to Alzheimer’s disease and related sex-disparities

**DOI:** 10.1101/321992

**Authors:** Alireza Nazarian, Anatoliy I. Yashin, Alexander M. Kulminski

**Author notes:** Corresponding Authors: Alireza Nazarian and Alexander M. Kulminski, Duke University, Bio-demography of Aging Research Unit, Social Science Research Institute, Erwin Mill Building, 2024 W. Main St., Durham, NC 27705. and.

## Abstract

**Background:** Alzheimer’s disease (AD) is the most common cause of dementia in the elderly and the sixth leading cause of death in the United States. AD is mainly considered a complex disorder with polygenic inheritance. Despite discovering many susceptibility loci, a major proportion of AD genetic variance remains to be explained.

**Methods:** We investigated the genetic architecture of AD in four publicly available independent datasets through genome-wide association, transcriptome-wide association, and gene-based analyses. To explore differences in the genetic basis of AD between males and females, analyses were performed on three samples in each dataset: males and females combined, only males, or only females.

**Results:** Our genome-wide association analyses corroborated the associations of several previously detected AD loci and revealed novel significant associations of 54 single-nucleotide polymorphisms (SNPs) at a p-value of < 5E-06. In addition, 23 genes located outside the chromosome 19q13 region showed suggestive associations with AD at a false discovery rate of 0.05 in transcriptome-wide association and gene-based analyses. Most of the newly detected AD-associated SNPs and genes were sex specific, indicating sex disparities in the genetic basis of AD.

**Conclusions:** Our findings, particularly the newly discovered sex-specific genetic contributors, provide novel insight into the genetic architecture of AD and can advance our understanding of its pathogenesis.

## Background

Alzheimer’s disease (AD) is a slowly progressive neurodegenerative disorder that usually manifests with insidious deterioration of cognitive functions such as memory, language, judgment, and reasoning. Visuospatial deficits and neuropsychiatric symptoms like anxiety, irritability, depression, delusion, and personality changes may occur in the course of the disease, and these are eventually followed by impairment of most daily activities [1,2]. The median survival is 3.3 to 11.7 years after disease manifestation [3]. Except for some uncommon autosomal dominant Mendelian forms, AD is mainly a complex disorder with a polygenic nature [2,4] that predominantly affects elderly individuals, also known as late-onset AD. It is the most common cause of dementia in elderly worldwide [5] and is the sixth leading cause of death in the United States [6]. Age is the main risk factor for AD. The annual incidence increases from 1&#x0025; at age 65 to 6–8&#x0025; after 85 years [7], and its prevalence increases from 11&#x0025; to 32&#x0025; [5]. In addition, AD is more prevalent and severe in females than males [7–10], with their lifetime risk of developing the disease being almost twice that of males [7]. The underlying mechanisms of sex disparity in AD are not clear [9,11]. This might be in part due to potential differences in the genetic bases of AD between males and females [12]. Investigating such differences is critical, particularly for tailoring more effective medical interventions [11,13].

Give the considerable physical, emotional, and economic burdens imposed by AD on patients, their families, and societies, exploring the genetic and non-genetic mechanisms underlying its pathogenesis has become a public health priority. With increased life expectancy, the prevalence and global economic costs of AD are forecast to increase considerably by 2050 [5]. Many studies have investigated the genetic basis of AD. Apolipoprotein E (APOE) was the first gene linked to late-onset AD [14], and, in particular, the dosage of its ε4 allele was implicated in increasing the risks of disease and earlier onset [15]. More susceptibility loci were detected with the advent of genome-wide association (GWA) methodology, although not all of them were consistently replicated in independent datasets. In addition to APOE, which was almost universally replicated, BIN1 [16–18], CLU [17–20], CR1 [17,19,20], CD2AP, CD33, MS4A4E, MS4A6A,EPHA1 [17,21], and PICALM [17–19] have been associated with the polygenic form of AD in different studies [22,23]. The narrow-sense heritability (h^2^) of AD (i.e., the proportion of its phenotypic variance explained by additive genetic variance) has been estimated to be 58–79&#x0025; by twin studies [24]. Furthermore, Ridge et al. (2016), using a linear mixed models (LMMs) framework, found out that 53&#x0025; of phenotypic variance of AD can be explained by ∼8 million single-nucleotide polymorphisms (SNPs). They also noticed that SNPs inside known AD-associated genes or within their 50 kb up-/downstream regions can only explaiñ31&#x0025; of AD phenotypic variance (∼59&#x0025; of genetic variance) [23], leaving a sizable portion of its h^2^ to be explained.

In this study, we investigated the genetic architecture of polygenic AD through genome-wide association(GWA), transcriptome-wide association, and gene-based analyses in four independent datasets (two with family designs and two with population designs) using genetic information for approximately two million genotyped and imputed SNPs. Since exploring the genetic sex disparity of AD was of particular interest, in addition to analyzing the entire sample of males and females in each dataset, two alternative plans were also considered in which either only males or only females were included in analyses.

## Methods

### Study Participants

Four independent datasets were used to fulfill the aims of this study: 1) Late-Onset Alzheimer’s Disease Family Study from the National Institute on Aging (NIA-LOADFS) [25], 2) Framingham SNP Health Association Resource (SHARe) project from Framingham Heart Study (FHS) [26–28], 3) SNP Typing for Association with Multiple Phenotypes from Existing Epidemiologic Data (STAMPEED) project from Cardiovascular Health Study (CHS) [29], and 4) the University of Michigan Health and Retirement Study (HRS) [30]. All four were approved by the institutional review boards (IRBs) and had gathered data after obtaining written informed consent from participants or their legal guardians/proxies. Details about the designs of NIA-LOADFS, FHS, CHS, and HRS studies can be found in the original publications. Our study focused on people of Caucasian ancestry from the four aforementioned studies to increase the sample size and power of the analyses. The LOADFS and FHS datasets directly identify cases with Alzheimer’s disease and unaffected controls. For the CHS and HRS datasets, International Classification of Disease codes, Ninth revision (ICD-9) were used to define cases and controls. Finally, to make the four datasets comparable in terms of participants age, we only included the original and offspring cohorts from the FHS dataset. Table S1 lists the numbers of cases and controls included in our study.

### Imputation of Genotype Data

Since the four datasets of interest were genotyped using different platforms, imputation was conducted to generate a common set of 2,928,658 SNPs. Only autosomal SNPs were subject to imputation. Genome coordinates of SNPs in our data (NCBI build 38/UCSC hg38) were lifted over to NCBI build 37/UCSC hg19 using liftover software [31]. After removing duplicate SNPs, pre-imputation quality control (QC) was performed using plink software [32] to remove low-quality SNPs/subjects by setting the following QC criteria: minor allele frequency < 0.01, SNPs and subject call rates < 95%, Hardy-Weinberg p-values < 1E-06. For the LOADFS and FHS cohorts that have family-based designs, a Mendel error rate of 2% was set to remove SNPs and subjects/families with high Mendelian errors. Strand alignment was checked using the SHAPEIT2 package [33] to ensure that allele calls are consistent between our and reference data. Haplotype phasing was then conducted using SHAPEIT2 to estimate the haplotypes for subjects in each dataset. Finally, genotypes were imputed by Minimac3 software [34] over pre-phased haplotypes. SHAPEIT2 and Minimac3 were run using default values for input arguments and European population (EUR) haplotypes from 1000 Genomes Phase 3 data (release October 2014) as the reference panel.

### Post-imputation QC

Directly genotyped SNPs along with the imputed SNPs, for which the squared correlation (r^2^) between imputed and expected true genotypes were above 0.7, were selected for pre-analysis QC. This step was performed based on the same criteria explained above for pre-imputation QC. Table S2 contains information on the numbers of genotyped and imputed SNPs that remained in each of the four datasets of interest after QC.

### Population Structure

The top 20 principal components (PCs) of genotype data were obtained through principal component analysis (PCA) to be included in downstream genetic analyses to address potential population stratification. In each dataset, PCA was performed over a subset of unrelated individuals and a subset of SNPs that were not in high linkage disequilibrium (LD) measured by r^2^ [35]. KING software [36] was used to obtain the subset of unrelated subjects by keeping one subject per family or relative cluster whose identity-by-descent (IBD) were above 0.0884 (i.e., closer than third-degree relatives). The genotyped autosomal SNPs on each chromosome were then pruned by plink software [32] in an unrelated set of subjects such that no SNP pairs with r^2^ > 0.2 were kept within any 100-SNPs windows. PCA was then conducted over the selected low-LD SNPs with the GENESIS R package [37]. Table S3 contains genomic inflation factors (λ values) resulted from logistic regression models for the four datasets under consideration. The λ values were less than 1.1 in all cases, indicating a subtle impact of population structure on our analyses [38,39].

### GWA Analysis

The associations between SNPs and AD were investigated by fitting logistic regression models. The genetic analyses of each dataset were performed under three alternative plans analyzing 1) the entire sample, 2) only males, and 3) only females. The top 5 PCs and subjects birth cohort were included in the models as fixed-effects covariates. In addition, sex was considered as a fixed effect covariate under plan 1. Only additive genetic effects were modeled; dominance effects were ignored. The logistic models were fitted using plink software [32]. It was previously suggested that for samples with family-based design, ignoring family relationships would not generate considerable bias in effect sizes of SNPs but may increase type I error rates whose magnitude depends on pedigree complexity (e.g., nuclear family vs. extended family) and trait heritability. For instance, the inflation of type I error rates has been suggested to be trivial in datasets with simple pedigrees. On the other hand, type I error rates may increase by a factor of two to three when family structure is ignored in a dataset with an extended family pedigree and trait heritability values of 0.6–0.9. Therefore, a two-step screening-validating approach could be used with such datasets to prevent inflation of type I error rates and decrease the computational burden of analysis [40]. For the LOADFS and FHS cohorts, we adopted a two-step approach in which the SNPs with p-values < 0.05 in the logistic models explained above were subjected to fitting generalized linear mixed models (GLMMs) by including all aforementioned fixed-effects covariates along with family IDs as a random effect covariate. GLMMs were fitted using lme4 R package [41].

All GWA analyses were conducted in a discovery-replication manner. Each of the LOADFS, FHS, CHS, and HRS datasets was considered as a discovery set to detect SNPs in significant associations with AD. Results from the discovery stage in a particular dataset were then subject to further replication in the remaining three datasets. At the discovery stage, a genome-wide significance level < 5E-08 was set to select statistically significant associations, and SNPs with p-values between 5E-08 and 5E-06 were considered as suggestive AD-associated markers. A significance threshold of 0.05 was considered during the replication stage.

Finally, a conventional fixed-effects meta-analysis, using the inverse variance method, was conducted over the results under each plan from the four investigated datasets to obtain combined statistics for the tested SNPs. To avoid missing heterogeneous associations of opposite directions of effects, we also performed a meta-analysis on absolute values of coefficients in addition to the conventional meta-test.

The results were interpreted according to the significance level at the discovery phase. The meta-analysis was performed using GWAMA software [42].

Significant findings from GWA analyses were compared to previous studies using the GRASP search tool [43]. Also, LD between significant SNPs and previously detected AD-associated loci in their 1 Mb flanking regions (r^2^ ≥ 0.4) was investigated in the CEU population (i.e., Utah Residents with Northern and Western European Ancestry) through the HaploR R package [44] and LDlink web-tool [45].

### Gene-based Analysis

Under each of three aforementioned plans, gene-based analysis was performed over the meta-analysis results using the fastBAT method [46] implemented in the GCTA package [47]. This method combines z-statistics of a set of SNPs corresponding to each gene into a quadratic form of a multivariate normal variable. SNPs located within a gene or its 50 kb up-/downstream regions were considered as an SNP set for that gene. The HRS dataset was used as the reference panel for LD calculation (i.e., r^2^ metric), and SNP pruning process in which one of each pair of SNPs with an r^2^ above 0.9 was removed from any given set. To deal with multiple-testing issue, the false discovery rate (FDR) method suggested by Benjamini and Hochberg was used to rank and select significant findings [48]. Genes with significant p-values at the FDR level of 0.05 were considered as novel AD-associated ones if there were no SNPs with P < 5E0−8 in their 1 Mb up-/downstream regions in the current or previous studies.

### Transcriptome-wide Association Analysis

Results from conducted meta-analyses along with summary data from a publicly available expression quantitative trait loci (eQTLs) study [49] were used to perform transcriptome-wide association analysis using SMR software [50]. The eQTLs summary data were downloaded from the SMR website. Both cis-and trans-eQTLs were of interest. Trans-eQTLs were defined as eQTLs located at least 5 Mb away from a probe on the same chromosome or located on other chromosomes. Probes for which at least one eQTL with P < 5E-08 had been detected by Lloyd-Jones et al. (2017) [49] were included in our analyses provided that the corresponding eQTLs were among the genotyped or imputed SNPs in our study. This resulted in the inclusion of sets of up to 8257 probes with cis-eQTLs and 2763 probes with trans-eQTLs.

The significance of p-values resulting from SMR testing (i.e., P_SMR_) were interpreted at an FDR level of 0.01 to 0.05. The appropriate FDR level for each of three analysis plans was chosen so we can ensure that the number of possible false-positive findings among significant probes was less than 1. As the aim of this analysis was to identify pleiotropic effects of SNPs on gene expression levels and AD development, probes with significant P_SMR_ were then subject to heterogeneity testing (i.e., the HEIDI test) which can differentiate pleiotropy (or causality) from linkage [50,51]. Genes corresponding to probes that passed both SMR and HEIDI tests (i.e., significant P_SMR_ and P_HEIDI_ ≥ 0.05) were deemed significant as their expression profiles might be associated with AD because of the pleiotropic effect of a single causal variant that affects both probe expression and AD susceptibility. Selected genes were considered potentially novel AD genes if there were no SNPs with P < 5E-08 within their 1 Mb up-/downstream regions in the current or previous studies.

### Availability of data and materials

The LOADFS, FHS, CHS, and HRS datasets are available through the dbGaP repository for qualified researchers (https://www.ncbi.nlm.nih.gov/gap).

## Results

### i. GWA Analysis

GWA analyses were performed in four independent datasets (i.e., LOADFS, FHS, CHS, and HRS). Each of these datasets served as discovery set to detect SNPs with significant association signals (at either a genome-wide significance level < 5E-08 or the suggestive level between 5E-08 and 5E-06), which were then subject to further replication (at significance level of 0.05) in the other three datasets. Finally, results from the individual datasets were combined through meta-analysis and interpreted according to the significance level at the discovery phase. Tables S4-S9 provide an overview of replicated and meta-analysis sets of SNPs that were significantly associated with AD in males and females combined (plan 1) or separately (plans 2 and 3). Figures S1-S6 show the Manhattan and QQ plots of the GWA results in the four investigated datasets, as well as in the conducted meta-analyses under these three plans. In general, SNPs with p-values smaller than the genome-wide significance threshold were mostly located on chromosome 19.

*Replicated sets of SNPs*: The replicated sets of SNPs under plans 1–3 contained 36, 26, and 30 SNPs, respectively (Tables S4-S6). These SNPs had significant p-values at the genome-wide level or suggestive level of associations at the discovery stage and were then replicated in another dataset. Notably, 14, 11, and 10 replicated SNPs, respectively, had not been previously associated with AD. Tables 1–3 summarize their information. The other SNPs had some evidence of direct association signals [43]. Among previously detected SNPs, rs9597722 (plan 1) [52], rs9882471 (plan 2) [53], and rs1359176 (plan 3) [54] were nominally associated with AD in previous studies (5E-06 ≤ P < 5E-02). Most of the newly detected SNPs were located inside a previously well-known susceptibility region for AD on chromosome 19q13 (i.e., APOE cluster genes region) and were mostly significant under different analysis plans. This subset of newly detected SNPs mostly had p-values less than 5E-08, the same directions of effects in discovery and replication datasets, and significant p-values (at genomic or suggestive level of significance) in meta-analysis. On the other hand, newly detected SNPs outside of the chromosome 19q13 region were mostly plan specific (except rs62402815). For example, SNPs that were significant in males were not significant in females. Also, they did not always have the same directions of effects in different datasets. Among these SNPs, rs62402815 under plans 1 and 3, and rs28385458 and rs726411 under plan 2 had the same direction of effects in multiple datasets. In addition, the association signals of this subset of newly detected SNPs were significant only at the suggestive level of associations (except rs62402815, which had a genome-wide level significant p-value of 1.2E-08 in females). Most of these SNPs did not have p-values smaller than 5E-06 in conventional fixed-effects meta-analyses, which might be partially due to the heterogeneity of their effects across different datasets. These heterogeneous effects were reflected by high i^2^ inconsistency metrics and significant Q-statistics in Cochran’s heterogeneity test (P_q_ < 0.05). A meta-analysis based on the absolute values of the coefficients confirmed a substantial role of heterogeneity by providing smaller p-values for most of these SNPs.

**Table 1:**
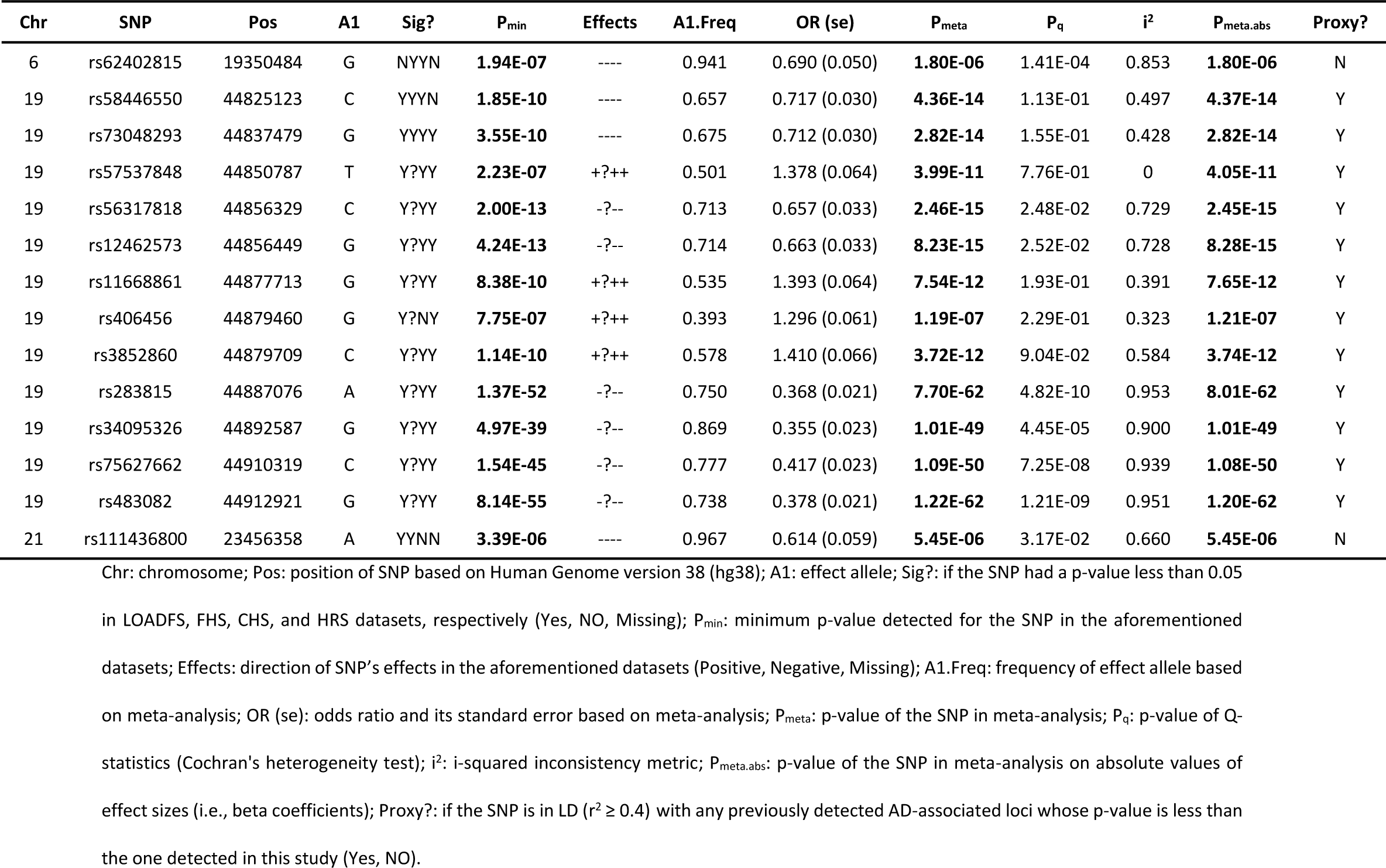
Newly detected replicated SNPs under plan 1 (males and females)

**Table 2:**
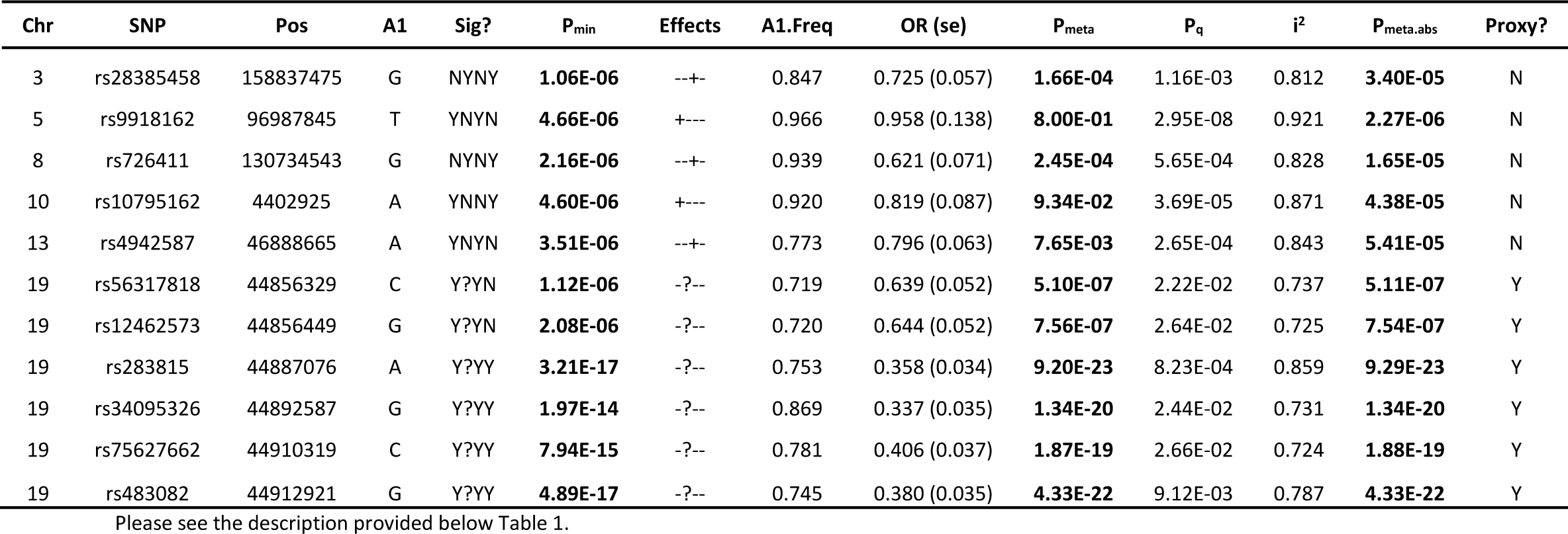
Newly detected replicated SNPs under plan 2 (only males)

**Table 3:** Newly detected replicated SNPs under plan 3 (only females)

| Chr | SNP | Pos | A1 | Sig? | P<sub>min</sub> | Effects | A1.Freq | OR (se) | P<sub>meta</sub> | P<sub>q</sub> | i<sup>2</sup> | P<sub>meta.abs</sub> | Proxy? |
| --- | --- | --- | --- | --- | --- | --- | --- | --- | --- | --- | --- | --- | --- |
| 6 | rs62402815 | 19350484 | G | NYYN | **1.20E-08** | ---- | 0.941 | 0.610 (0.054) | **4.29E-07** | 5.89E-04 | 0.827 | **4.31E-07** | N |
| 19 | rs58446550 | 44825123 | C | YYNY | **3.96E-06** | ---- | 0.653 | 0.712 (0.038) | **1.21E-09** | 6.49E-01 | 0 | **1.21E-09** | Y |
| 19 | rs56317818 | 44856329 | C | Y?YY | **7.66E-08** | -?-- | 0.710 | 0.663 (0.042) | **1.24E-09** | 2.42E-01 | 0.295 | **1.23E-09** | Y |
| 19 | rs12462573 | 44856449 | G | Y?NY | **8.15E-08** | -?-- | 0.710 | 0.669 (0.042) | **2.72E-09** | 2.08E-01 | 0.363 | **2.72E-09** | Y |
| 19 | rs11668861 | 44877713 | G | Y?NY | **1.93E-07** | +?++ | 0.532 | 1.376 (0.079) | **1.87E-07** | 7.94E-02 | 0.605 | **1.86E-07** | Y |
| 19 | rs35879138 | 44879882 | T | Y?YN | **8.45E-07** | -?-- | 0.892 | 0.644 (0.056) | **3.44E-06** | 3.33E-02 | 0.706 | **3.44E-06** | Y |
| 19 | rs283815 | 44887076 | A | Y?YY | **3.72E-31** | -?-- | 0.748 | 0.384 (0.028) | **3.17E-35** | 1.05E-06 | 0.927 | **3.14E-35** | Y |
| 19 | rs34095326 | 44892587 | G | Y?YY | **4.83E-23** | -?-- | 0.869 | 0.369 (0.031) | **3.81E-27** | 3.01E-04 | 0.877 | **3.86E-27** | Y |
| 19 | rs75627662 | 44910319 | C | Y?YY | **1.92E-29** | -?-- | 0.773 | 0.424 (0.030) | **1.02E-29** | 7.43E-07 | 0.929 | **1.02E-29** | Y |
| 19 | rs483082 | 44912921 | G | Y?YY | **2.45E-35** | -?-- | 0.734 | 0.382 (0.027) | **1.21E-37** | 3.97E-08 | 0.941 | **1.20E-37** | Y |

Please see the description provided below Table 1.

Although no evidence of direct association with AD was found in previous studies for the newly detected subsets of replicated SNPs, their 1 Mb up-/downstream regions harbor AD-associated SNPs. We therefore investigated their LD with AD-associated loci in their 1 Mb flanking regions in the CEU population [45]. Newly detected SNPs were considered novel AD markers if their p-values were smaller than those of the top AD-associated SNPs in their neighborhood or they were not in LD (r^2^ ≥ 0.4) with previously AD-associated loci whose p-values were smaller than those detected in this study. SNPs on chromosome 19q13 mostly had larger p-values than top AD-associated loci in their neighborhood and were in LD with them. On the other hand, the p-values of SNPs located outside chromosome 19q13 region were mostly smaller than the previously detected association signals in their flanking regions and were not in LD with such loci. Table S10 contains LD information about those newly detected SNPs for which proxy AD-associated loci have been reported.

*Meta-analysis sets of SNPs*: Tables S7-S9 show that 17, 4, and 24 SNPs that were not among replicated sets of SNPs under analysis plans 1–3 passed the significance threshold in meta-analysis. The meta-analysis p-values of these SNPs were mostly significant at the level of suggestive associations, except for rs57537848 and rs76366838 on chromosome 19q13 with P < 5E-08 in females. Also, they were mostly located outside chromosome 19q13 and were plan specific. For example, significant SNPs in males were not significant in females and vice versa. In addition, most (14, 3, and 24 SNPS under plans 1–3, respectively) were not associated with AD in previous studies [43]. Summary information about the newly detected subset of meta-analysis sets of SNPs can be found in Tables 4–6. As with replicated sets of SNPs, the newly detected SNPs not on chromosome 19q13 either had smaller p-values than the ones reported for their nearby AD-associated loci or were not in LD with their corresponding AD-associated SNPs (Table S10).

**Table 4:**
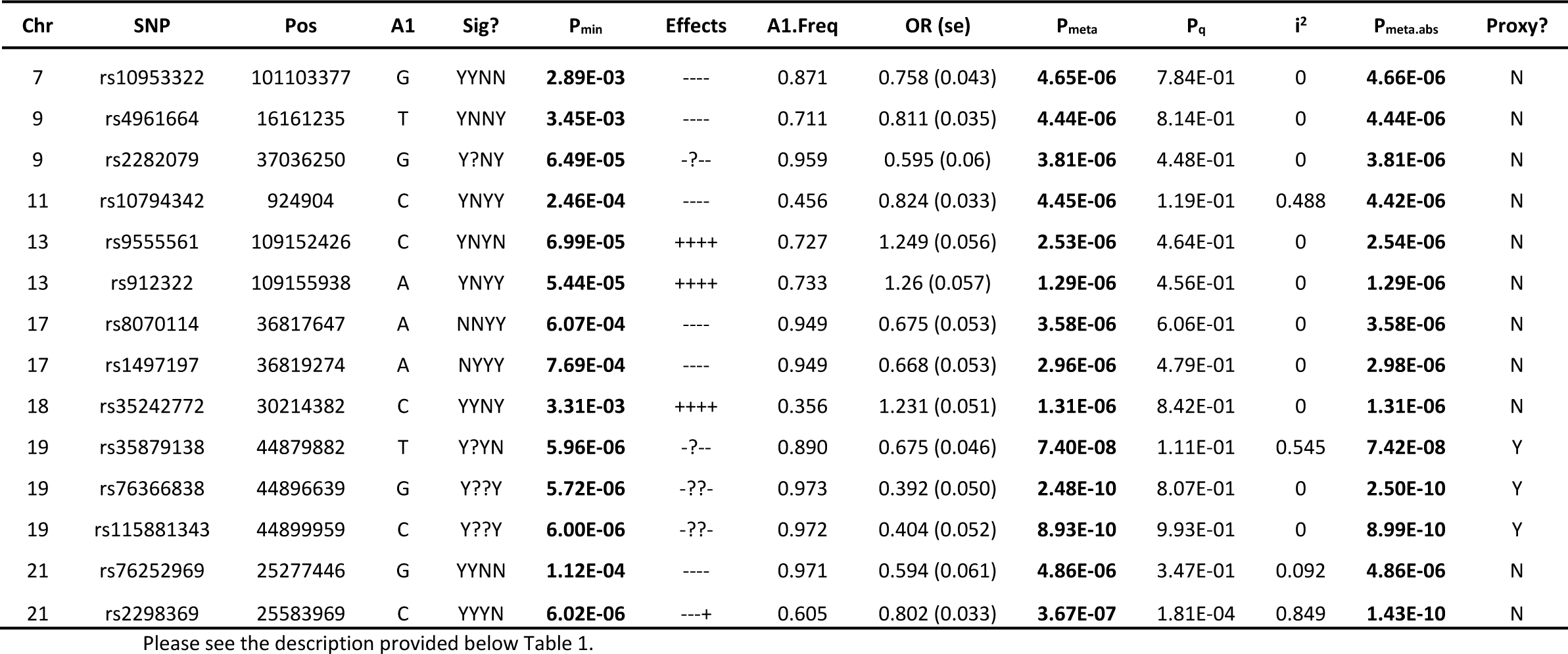
Newly detected meta-analysis set of SNPs under plan 1 (males and females)

**Table 5:** Newly detected meta-analysis set of SNPs under plan 2 (only males)

| Chr | SNP | Pos | A1 | Sig? | P<sub>min</sub> | Effects | A1.Freq | OR (se) | P<sub>meta</sub> | P<sub>q</sub> | i<sup>2</sup> | P<sub>meta.abs</sub> | Proxy? |
| --- | --- | --- | --- | --- | --- | --- | --- | --- | --- | --- | --- | --- | --- |
| 3 | rs9862849 | 66855351 | C | YNNY | **1.12E-03** | ---- | 0.900 | 0.603 (0.059) | **2.66E-06** | 7.53E-01 | 0 | **2.65E-06** | N |
| 19 | rs3852860 | 44879709 | C | Y?YY | **1.36E-03** | +?++ | 0.581 | 1.480 (0.115) | **3.53E-06** | 9.28E-01 | 0 | **3.55E-06** | Y |
| 23 | rs5969117 | 87181248 | C | YN?N | **1.67E-05** | ++?+ | 0.295 | 1.490 (0.114) | **1.42E-06** | 8.61E-01 | 0 | **1.42E-06** | N |

Please see the description provided below Table 1.

**Table 6:**
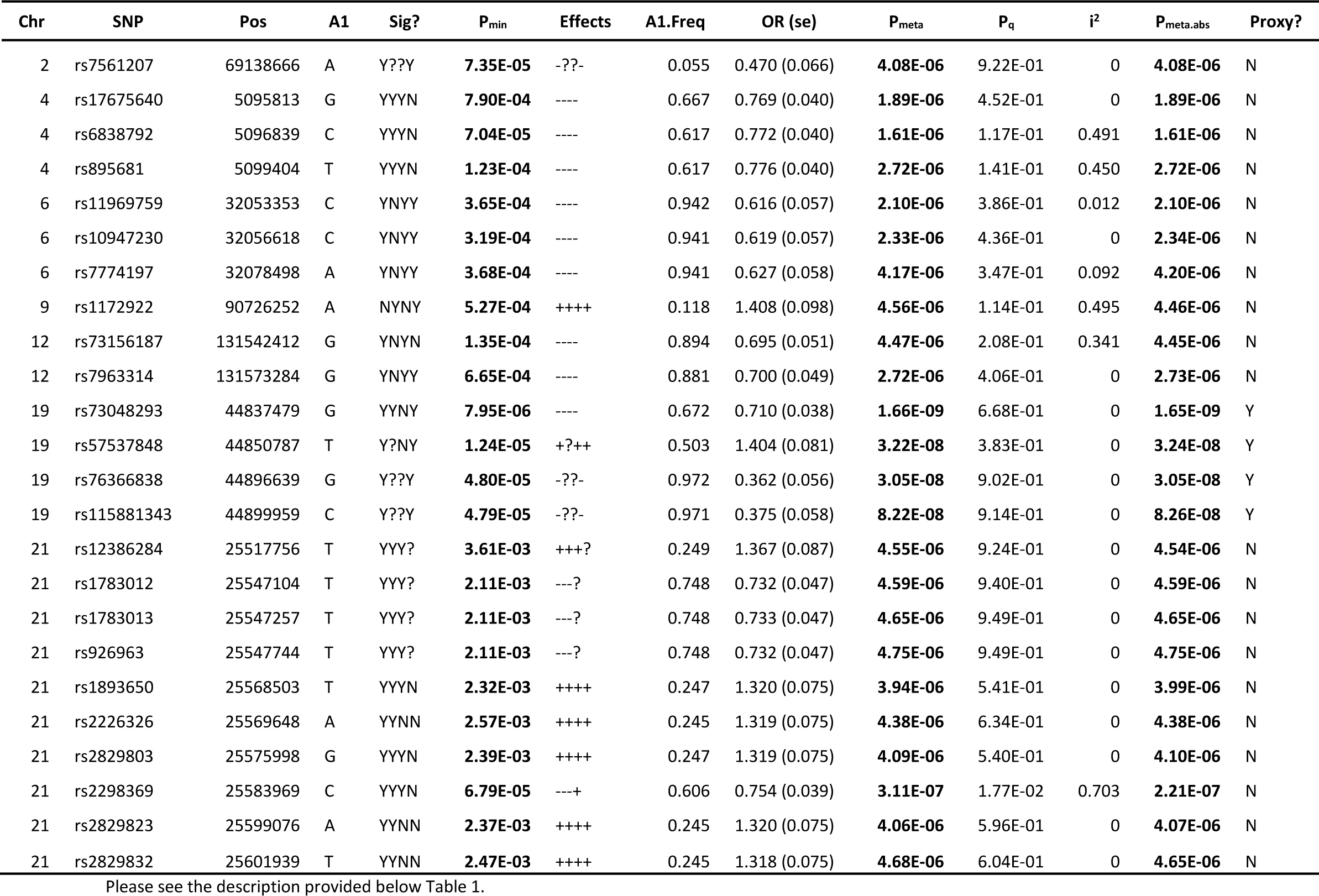
Newly detected meta-analysis set of SNPs under plan 3 (only females)

### ii. Gene-based Analysis

The significant findings from gene-based analyses corresponding to plans 1–3 are summarized in Table 7. Under all plans, most genes with significant p-values at the FDR of 0.05 were located in the chromosome 19q13 region. Since chromosome 19q13 region harbors several SNPs with p-values less than 5E-08 in both current and previous studies, significant genes in this region are not discussed here as they do not meet the criteria set for detecting novel AD genes. The only significant genes outside this APOE cluster region were LINC00158 under plan 1 and LINC00158, MIR155HG, MIR155, LINC00515, MRPL39, and JAM2 genes under plan 3. None of the SNPs inside or in 1 Mb flanking regions of these genes had significant p-values at the genome-wide level in our study, although several had suggestive level p-values in conducted meta-analyses under plans 1 and 3. Also, SNPs in their nearby regions were only nominally associated with AD (8.0E-04 < P < 5E-02) in previous studies [53–57].

**Table 7:**
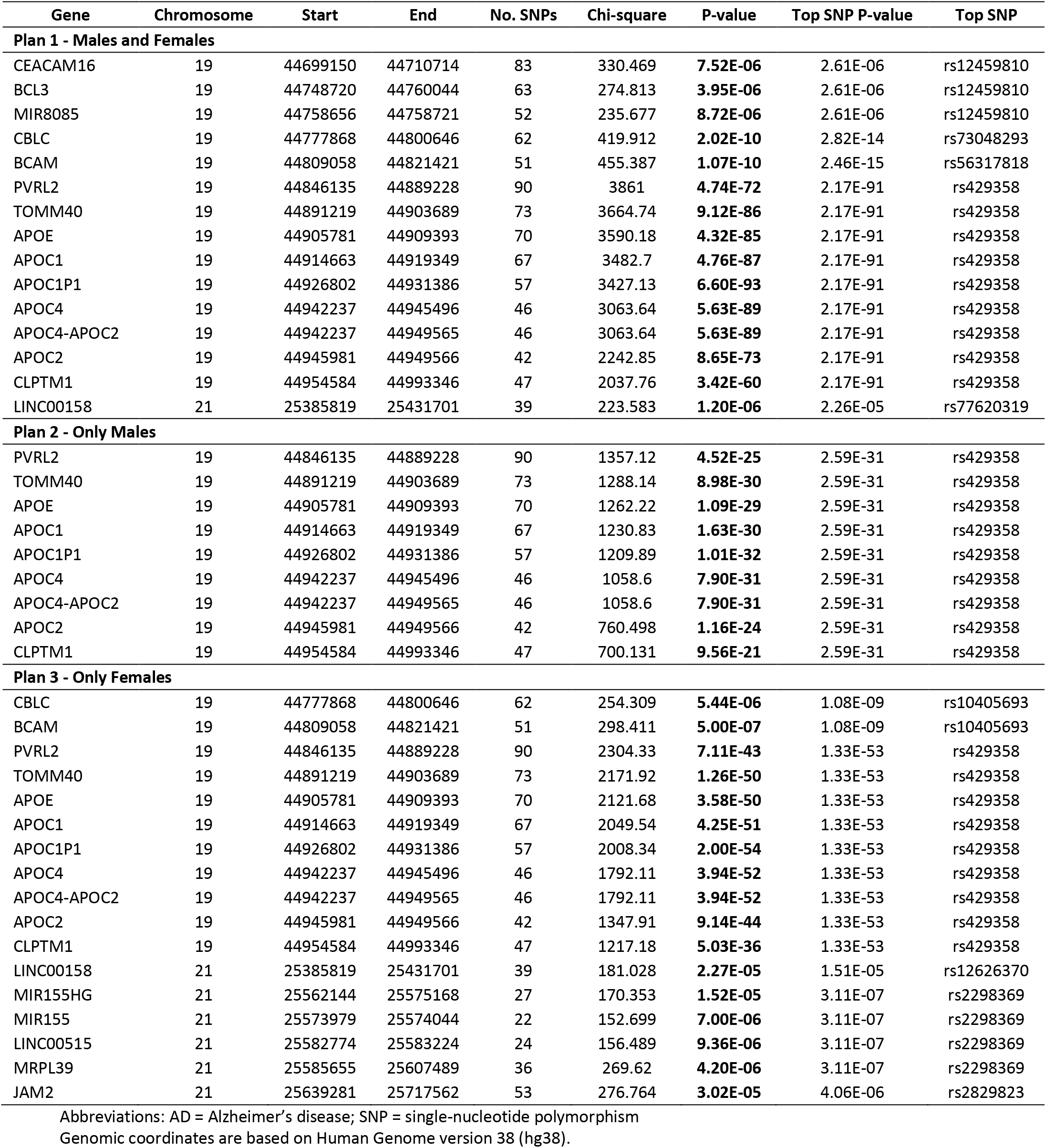
Significantly AD-associated genes from gene-based analyses.

| Gene | Chromosome | Start | End | No. SNPs | Chi-square | P-value | Top SNP P-value | Top SNP |
| --- | --- | --- | --- | --- | --- | --- | --- | --- |
| **Plan 1 - Males and Females** |  |  |  |  |  |  |  |  |
| CEACAM16 | 19 | 44699150 | 44710714 | 83 | 330.469 | **7.52E-06** | 2.61E-06 | rs12459810 |
| BCL3 | 19 | 44748720 | 44760044 | 63 | 274.813 | **3.95E-06** | 2.61E-06 | rs12459810 |
| MIR8085 | 19 | 44758656 | 44758721 | 52 | 235.677 | **8.72E-06** | 2.61E-06 | rs12459810 |
| CBLC | 19 | 44777868 | 44800646 | 62 | 419.912 | **2.02E-10** | 2.82E-14 | rs73048293 |
| BCAM | 19 | 44809058 | 44821421 | 51 | 455.387 | **1.07E-10** | 2.46E-15 | rs56317818 |
| PVRL2 | 19 | 44846135 | 44889228 | 90 | 3861 | **4.74E-72** | 2.17E-91 | rs429358 |
| TOMM40 | 19 | 44891219 | 44903689 | 73 | 3664.74 | **9.12E-86** | 2.17E-91 | rs429358 |
| APOE | 19 | 44905781 | 44909393 | 70 | 3590.18 | **4.32E-85** | 2.17E-91 | rs429358 |
| APOC1 | 19 | 44914663 | 44919349 | 67 | 3482.7 | **4.76E-87** | 2.17E-91 | rs429358 |
| APOC1P1 | 19 | 44926802 | 44931386 | 57 | 3427.13 | **6.60E-93** | 2.17E-91 | rs429358 |
| APOC4 | 19 | 44942237 | 44945496 | 46 | 3063.64 | **5.63E-89** | 2.17E-91 | rs429358 |
| APOC4-APOC2 | 19 | 44942237 | 44949565 | 46 | 3063.64 | **5.63E-89** | 2.17E-91 | rs429358 |
| APOC2 | 19 | 44945981 | 44949566 | 42 | 2242.85 | **8.65E-73** | 2.17E-91 | rs429358 |
| CLPTM1 | 19 | 44954584 | 44993346 | 47 | 2037.76 | **3.42E-60** | 2.17E-91 | rs429358 |
| LINC00158 | 21 | 25385819 | 25431701 | 39 | 223.583 | **1.20E-06** | 2.26E-05 | rs77620319 |
| **Plan 2 - Only Males** |  |  |  |  |  |  |  |  |
| PVRL2 | 19 | 44846135 | 44889228 | 90 | 1357.12 | **4.52E-25** | 2.59E-31 | rs429358 |
| TOMM40 | 19 | 44891219 | 44903689 | 73 | 1288.14 | **8.98E-30** | 2.59E-31 | rs429358 |
| APOE | 19 | 44905781 | 44909393 | 70 | 1262.22 | **1.09E-29** | 2.59E-31 | rs429358 |
| APOC1 | 19 | 44914663 | 44919349 | 67 | 1230.83 | **1.63E-30** | 2.59E-31 | rs429358 |
| APOC1P1 | 19 | 44926802 | 44931386 | 57 | 1209.89 | **1.01E-32** | 2.59E-31 | rs429358 |
| APOC4 | 19 | 44942237 | 44945496 | 46 | 1058.6 | **7.90E-31** | 2.59E-31 | rs429358 |
| APOC4-APOC2 | 19 | 44942237 | 44949565 | 46 | 1058.6 | **7.90E-31** | 2.59E-31 | rs429358 |
| APOC2 | 19 | 44945981 | 44949566 | 42 | 760.498 | **1.16E-24** | 2.59E-31 | rs429358 |
| CLPTM1 | 19 | 44954584 | 44993346 | 47 | 700.131 | **9.56E-21** | 2.59E-31 | rs429358 |
| **Plan 3 - Only Females** |  |  |  |  |  |  |  |  |
| CBLC | 19 | 44777868 | 44800646 | 62 | 254.309 | **5.44E-06** | 1.08E-09 | rs10405693 |
| BCAM | 19 | 44809058 | 44821421 | 51 | 298.411 | **5.00E-07** | 1.08E-09 | rs10405693 |
| PVRL2 | 19 | 44846135 | 44889228 | 90 | 2304.33 | **7.11E-43** | 1.33E-53 | rs429358 |
| TOMM40 | 19 | 44891219 | 44903689 | 73 | 2171.92 | **1.26E-50** | 1.33E-53 | rs429358 |
| APOE | 19 | 44905781 | 44909393 | 70 | 2121.68 | **3.58E-50** | 1.33E-53 | rs429358 |
| APOC1 | 19 | 44914663 | 44919349 | 67 | 2049.54 | **4.25E-51** | 1.33E-53 | rs429358 |
| APOC1P1 | 19 | 44926802 | 44931386 | 57 | 2008.34 | **2.00E-54** | 1.33E-53 | rs429358 |
| APOC4 | 19 | 44942237 | 44945496 | 46 | 1792.11 | **3.94E-52** | 1.33E-53 | rs429358 |
| APOC4-APOC2 | 19 | 44942237 | 44949565 | 46 | 1792.11 | **3.94E-52** | 1.33E-53 | rs429358 |
| APOC2 | 19 | 44945981 | 44949566 | 42 | 1347.91 | **9.14E-44** | 1.33E-53 | rs429358 |
| CLPTM1 | 19 | 44954584 | 44993346 | 47 | 1217.18 | **5.03E-36** | 1.33E-53 | rs429358 |
| LINC00158 | 21 | 25385819 | 25431701 | 39 | 181.028 | **2.27E-05** | 1.51E-05 | rs12626370 |
| MIR155HG | 21 | 25562144 | 25575168 | 27 | 170.353 | **1.52E-05** | 3.11E-07 | rs2298369 |
| MIR155 | 21 | 25573979 | 25574044 | 22 | 152.699 | **7.00E-06** | 3.11E-07 | rs2298369 |
| LINC00515 | 21 | 25582774 | 25583224 | 24 | 156.489 | **9.36E-06** | 3.11E-07 | rs2298369 |
| MRPL39 | 21 | 25585655 | 25607489 | 36 | 269.62 | **4.20E-06** | 3.11E-07 | rs2298369 |
| JAM2 | 21 | 25639281 | 25717562 | 53 | 276.764 | **3.02E-05** | 4.06E-06 | rs2829823 |

Abbreviations: AD = Alzheimer's disease; SNP = single-nucleotide polymorphism Genomic coordinates are based on Human Genome version 38 (hg38).

### iii. Transcriptome-wide Association Analysis

*a. Analyzing probes with cis-eQTLs:* We found that 10, 17, and 15 probes/genes had significant p-values in SMR test under plans 1–3, respectively. The significant FDR level for interpreting the results from SMR test was set to 0.05 under plan 1 and 0.025 under plans 2 and 3 to ensure that the expected number of false-positive findings was less than 1. Of these, only 4, 8, and 4 probes/genes (with P_SMR_< 6.03E-05) passed heterogeneity testing (P_HEIDI_ ≥ 0.05). Table 8 contains information about these 16 probes/genes, their top eQTLs, and respective p-values. The top eQTLs corresponding to these probes/genes were all nominally significant in our GWA analyses (2.01E-04 ≤ P_GWAS_ ≤ 2.47E-02). Moreover, we did not identify any SNPs with significant p-values at the genome-wide significance level within 1 Mb of these genes. However, several SNPs within 1 Mb of MS4A6A [17,21,58,59] and UQCC [60] were associated with AD with P < 5E-08 in previous studies. Among 14 other genes, SNPs in regions around TRA2A [59], IRAK3 [61], and ESPN [62] were previously associated with AD at the suggestive level of associations.

**Table 8:**
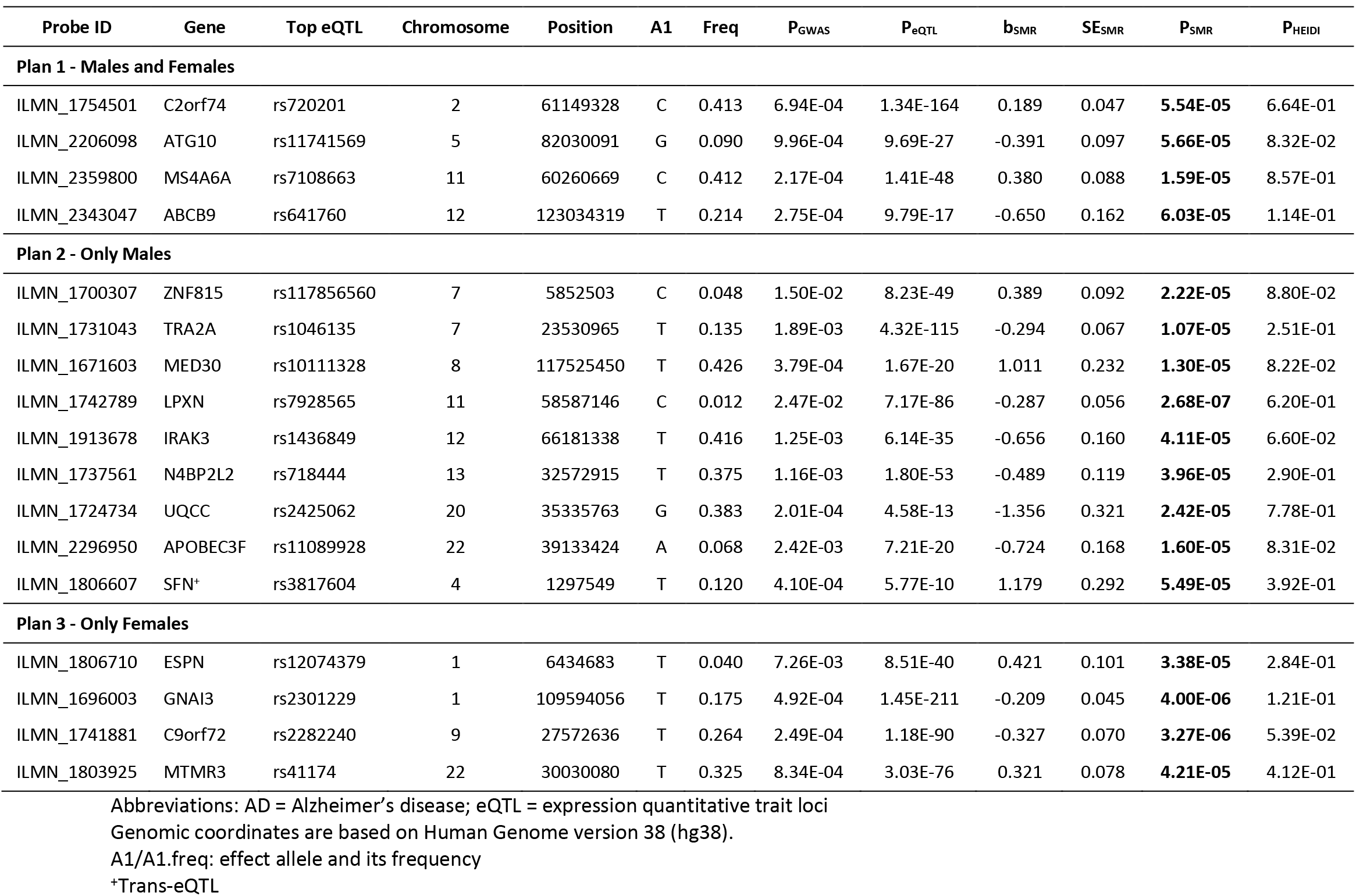
Significantly AD-associated probes/genes from transcriptome-wide analyses.

*b. Analyzing probes with trans-eQTLs:* No probe passed both SMR and HEIDI tests under analysis plans 1 and 3. However, 7 probes had significant P_SMR_ at the FDR level of 0.05 under plan 2, of which only one probe mapping to SFN gene on chromosome 1p36 passed the HEIDI test (Table 8). The corresponding top eQTL was located on chromosome 4p16 in the intronic region of the MAEA gene and was nominally associated with AD in our study (P_GWAS_ = 4.10E-04). There were no significant association signals at the genome-wide significance level in the SFN gene or its 1 Mb flanking regions in current or previous studies.

## Discussion

The genetic architecture of AD has been widely studied in recent years, and so far more than 60000 SNPs have been associated with AD with P < 0.05. Of these, 281 SNPs (mapped to 49 genes) and 593 SNPs (mapped to 165 genes) had significant p-values at the genome-wide and suggestive levels of associations, respectively [43]. Despite these efforts, a major proportion of h^2^ of AD has remained unexplained. Exploring the genetic risk factors contributing to AD is highly important from a precision medicine perspective where the goal is to personalize diagnostic and therapeutic interventions. The current study provides further insight into the genetic architecture of AD through GWA, transcriptome-wide association, and gene-based analyses of four independent datasets.

Our GWA analyses corroborated the associations of a number of previously detected AD loci and revealed some significant novel association signals. Among previously detected AD-associated SNPs, we found several SNPs with p-values that were smaller than those reported before. Also, the significant association signals for two SNPs inside chromosome 19q13 region (i.e., rs10426423 from the meta-analysis set of SNPs under plan 1, and rs769450 from the replicated sets of SNPs under plans 1 and 3) were previously reported only in African-Americans (p-values of 2.6E-8, 9.9E-7, and 5.3E-27, respectively [63]). Most newly detected AD-associated SNPs, particularly those outside chromosome 19q13 region, can be considered as novel AD markers because their p-values in our study were smaller than those for other AD-associated loci in their 1 Mb up-/downstream regions or they are not in LD with such loci. For instance, as seen in Tables 1–6 that summarize the replicated and meta-analysis sets of SNPs, 13, 9, and 21 novel AD-associated SNPs were detected under plans 1–3, respectively.

In the gene-based analysis, LINC00158, MIR155HG, MIR155, LINC00515, MRPL39, and JAM2 genes were significantly associated with AD when the entire sample of individuals and/or only females were analyzed. These genes are located near each other on chromosome 21q21 in a ∼332 kb region. The APP gene implicated in early onset familial AD is also located 163–449 kb from these genes. There were no AD-associated SNPs with p-values smaller than 5E-08 within their 1Mb in current or previous studies [43]. However, there were several SNPs with significant p-values at the suggestive level of associations in that chromosomal region among meta-analysis sets of SNPs under plans 1 and 3. The SNPs in the 1 Mb up/downstream regions of these genes were previously associated with some potential AD risk factors such as type 2 diabetes, hypertension, coronary artery disease, and lipid profile changes at the genome-wide significance level. They have also been associated with traits such as alcohol and nicotine co-dependence, age at onset of Parkinson’s disease, and pattern recognition memory at the suggestive significance level of association [43]. Furthermore, functional studies have provided insight into the potential roles of some of these genes in AD pathogenesis. LINC00158 and LINC00515 encode two long intergenic non-coding RNAs, and MIR155HG and MIR155 encode two microRNAs. MIR155 overexpression was previously implicated in downregulation of complement factor H (CFH) expression in AD and other neurodegenerative diseases. CFH encodes a regulatory protein that prevents spontaneous immune system activation [64]. MRPL39 encodes a mitochondrial ribosomal protein involved in the oxidative-phosphorylation pathway. Impaired mitochondrial function has been reported in neurons of patients with AD [65,66]. Lunnon et al. (2017) reported that the expression levels of MRPL39 and another nearby gene (i.e., ATP5J involved in the oxidative-phosphorylation pathway) were slightly reduced in AD patients compared to controls [65]. JAM2 encodes a membrane protein found at the tight junctions of epithelial and endothelial cells that acts as an adhesive ligand for immune cells. It belongs to the immunoglobulin super-family of adhesive molecules that has been implicated in AD pathogenesis [67]. Also, duplication of a ∼600 kb region on chromosome 21 containing the JAM2, ATP5J, and APP genes has been reported in autosomal dominant AD [68].

In transcriptome-wide association analyses, the expression level of 17 probes/genes passed both SMR and HEIDI tests, indicating that causal variants influencing the expression of these genes may also have pleiotropic effects on developing AD [50,51]. It should be noted that although using data from eQTLs studies on blood is not ideal for capturing associations between the brain-specific transcriptome and AD, however, it increases the power of SMR analysis since such studies take advantage of more samples compared to tissue-specific eQTLs studies [50]. Except for MS4A6A and UQCC genes, no significant SNPs with P < 5E-08 were detected within 1 Mb of the other genes in our GWAS or previous reports [43], although several SNPs with P < 5E-06 were reported in regions around TRA2A [59], IRAK3 [61], and ESPN [62]. This is likely due to the insufficient sample sizes and, consequently, the lack of power of conducted GWAS [50]. Taken together, all these genes except MS4A6A and UQCC can be considered as novel potential AD-associated genes. Further follow-up functional analyses are needed to explore their potential roles in AD pathogenesis. Among significant genes detected in females, a pathologic hexa-nucleotide repeat expansion in the C9orf72 gene has been linked to frontotemporal dementia and may contribute to AD pathogenesis [69–72]. Also, the GNAI3 gene was reported to be overexpressed in AD intact mice compared to AD impaired ones [73]. Moreover, SNPs in 1 Mb up-/downstream regions around these genes have been previously associated with some other traits (e.g., autoimmune diseases or serum cholesterol levels) with P < 5E-06. Examples include associations of SNPs corresponding to ATG10 with vascular dementia, ABCB9 with college completion and years of education, MED30 with rheumatoid arthritis and fasting blood glucose, LPXN with inflammatory bowel disease, GNAI3 with total and low-density lipoprotein cholesterol (LDL) and major depression, and SFN with high-density lipoprotein cholesterol (HDL) [43].

Finally, of particular interest was to investigate the sex disparity in the genetic basis of AD. Addressing sex differences in biomedical research has been emphasized by the National Institutes of Health as an approach that can eventually bolster the personalized medicine paradigm [11,13]. Our results revealed a number of new sex-specific genetic contributors to AD at SNP, gene, and transcriptome levels. For instance, most of the newly detected SNPs, particularly SNPs outside of chromosome 19q13, were sex specific. Another level of sex disparity was observed in the gene-based and transcriptome-wide association analyses. Notably, none of the novel AD-associated genes detected in males were among significant genes in females and vice versa.

In summary, our study revealed significant associations of 54 novel SNPs at suggestive or genomic levels of significance (Tables 1–6). Most (39) of these SNPs were located outside of 19q13 (i.e., APOE cluster genes region) and were not in LD (r^2^ ≥ 0.4) with SNPs previously associated with AD with P < 5E-06. Also, 23 genes located outside the chromosome 19q13 region showed evidence of association with AD at the FDR level of 0.05 in our transcriptome-wide association and gene-based analyses. Most of these AD-associated SNPs and genes were sex specific, indicating sex disparities in the genetic basis of AD. By detecting a number of novel AD-associated SNPs and discovering suggestive associations of several genes and transcripts, our study provides new insight into the genetic architecture of AD. Particularly, identifying sex-specific genetic contributors can advance our understanding of AD pathogenesis. Further studies, possibly with larger sample sizes, are needed to clarify the genotype-phenotype relationships in AD.

## Acknowledgments

This research was supported by Grants from the National Institute on Aging (P01AG043352 and R01AG047310). The funders had no role in study design, data collection and analysis, decision to publish, or manuscript preparation. The content is solely the responsibility of the authors and does not necessarily represent the official views of the National Institutes of Health.

This manuscript was prepared using limited access datasets obtained though dbGaP (accession numbers: phs000168.v2.p2 (LOADFS), phs000007.v28.p10 (FHS), phs000287.v5.p1 (CHS), and phs000428.v2.p2 (HRS)) and the University of Michigan. Phenotypic HRS data are available publicly and through restricted access from http://hrsonline.isr.umich.edu/index.php?p&#x003D;data. The authors thank Arseniy P. Yashkin for help preparing the HRS phenotypes. See also Supplementary Acknowledgment File.

## Supplementary information

Supplementary File 1 and Supplementary Acknowledgment File.

